# Phages actively challenge niche communities in the Antarctic soils

**DOI:** 10.1101/2020.01.17.911081

**Authors:** Oliver K.I Bezuidt, Pedro Humberto Lebre, Rian Pierneef, Carlos León-Sobrino, Evelien M. Adriaenssens, Don A. Cowan, Yves Van de Peer, Thulani P. Makhalanyane

**Author notes:** Oliver K.I. Bezuidt and Pedro H Lebre contributed equally to this work. Author order was determined on the basis of seniority. Address correspondence to Thulani P. Makhalanyane.

## Abstract

By modulating the structure, diversity and trophic outputs of microbial communities, phages play crucial roles in many biomes. In oligotrophic polar deserts, the effects of katabatic winds, constrained nutrients and low water availability are known to limit microbial activity. Although phages may substantially govern trophic interactions in cold deserts, relatively little is known regarding the precise ecological mechanisms. Here, we provide the first evidence of widespread antiphage innate immunity in Antarctic environments using metagenomic sequence data from hypolith communities as model systems. In particular, immunity systems such as DISARM and BREX are shown to be dominant systems in these communities. Additionally, we show a direct correlation between the CRISPR-cas adaptive immunity and the metavirome of hypolith communities, suggesting the existence of dynamic host-phage interactions. In addition to providing the first exploration of immune systems in cold deserts, our results suggest that phages actively challenge niche communities in Antarctic polar deserts. We provide evidence suggesting that the regulatory role played by phages in this system is an important determinant of bacterial host interactions in this environment.

**Importance:** In Antarctic environments, the combination of both abiotic and biotic stressors results in simple trophic levels dominated by microbiomes. Although the past two decades have revealed substantial insights regarding the diversity and structure of microbiomes, we lack mechanistic insights regarding community interactions and how phages may affect these. By providing the first evidence of widespread antiphage innate immunity, we shed light on phage-host dynamics in Antarctic niche communities. Our analyses reveal several antiphage defense systems including DISARM and BREX, which appear to dominate in cold desert niche communities. In contrast, our analyses revealed that genes, which encode antiphage adaptive immunity were under-represented in these communities suggesting lower infection frequencies in cold edaphic environments. We propose that by actively challenging niche communities, phages play crucial roles in the diversification of Antarctic communities.

## Introduction

Antarctic terrestrial environments including open soils, permafrost and the surface and interior of rocks, are typically oligotrophic and dominated by psychrophilic and psychrotolerant microbial communities (1–4). It has been suggested that the extreme abiotic pressures of the environment such as temperature, desiccation stress and UV radiation are dominant drivers of both the diversity and function of cold-adapted bacterial communities in terrestrial polar deserts (5–7). Similarly, biotic interactions such as competition, symbioses, horizontal gene transfer (HGT) and predation have also been shown to play a role in the distribution and diversity of microbial communities in these soil ecosystems (8–10). The presence of viruses, including bacteriophages, in these cold hyper-arid desert soils potentially adds an additional layer of complexity to the microbial system, but the extent to which phage-host interactions play a role in shaping community compositions and processes in cold desert soil niches remains a matter of speculation (11, 12).

Antarctic desert hypolithic communities, in particular, have been shown to contain substantial viral populations, dominated by tailed bacteriophages of the order *Caudovirales* (11, 13-15). Micro-array analysis of lithic niches identified an even greater phage diversity, including signatures of RNA bacteriophages of the family *Leviviridae,* ssDNA phage of the family *Microviridae* and non-tailed dsDNA tectiviruses (16). Together, these observations suggest that phages may play an important role of in microbial community structures and functions.

The presence of active bacteriophages in a microbial community inevitably leads to the evolution of specialized bacterial defensive measures (17), and a diverse range of bacterial defense mechanisms against parasitic phages have been identified (18, 19). These include adaptive immunity elements, such as the CRISPR-Cas systems, and innate immunity mechanisms, such as restriction-modification (RM) and toxin-antitoxin abortive infection (Abi) systems (18). Recent pangenomics studies have also identified novel defense systems that are widely distributed across bacterial taxa and are thought to play a role in anti-phage resistance (20–23). These include the bacteriophage exclusion (BREX) system, coded by a 4-8 gene cluster, that provides resistance to *Siphoviridae* and *Myoviridae* tailed phages by inhibition of phage DNA replication (21), and other less well characterized systems such as the Thoeris, Shedu and Gabija elements that increase bacterial host resistance to specific groups of phages (22).

Combining the valuable evidence on phage diversity and prevalence in polar desert soils, we hypothesize that phage-host interactions play an important role in shaping the structure of edaphic microbial communities in these environments. To test our hypothesis, we assess the known bacterial defense systems in metagenomic sequence data derived from niche Antarctic hypolith community. We were able to link some of these data to specific phage genomes and propose that phages play an active role in shaping the immunity of Antarctic soil microbial communities.

## Results

### The distribution of anti-phage defense mechanisms shows an abundance of innate immunity genes

The distribution of antiphage defense systems in the metagenome was determined by mapping defense genes against the taxonomically assigned contigs. In total, 24,941 defense genes were detected, compromising 1.2% of the entire metagenome gene count. Approximately 40% of these were found in contigs attributed to unknown phyla. The general distribution of defense genes across known phyla was consistent with the relative abundance of each phylum in the metagenome (Figure 1A, Table S4). Proteobacteria harbored the highest number of anti-phage genes (5289 genes, 1.1% of total gene count for this phylum), followed by Actinobacteria (3808, 0.9% total gene count) and Bacteroidetes (2128, 1.08% of total contig count). RM, DISARM and BREX systems were the most abundant systems in the metagenome, contributing 67.6% of the total gene hits for anti-phage defense systems. On the other side of the spectrum, the defense systems Shedu, Hachiman and CRISPR-type 2 were present at relatively low abundances, and therefore had little apparent contribution to the global defense system distribution. The average contribution of defense genes to the total gene count per phyla was 1.8%, with Deferribacteres and Candidatus Tectomicrobia as outliers. However, it is important to note that these phyla represent a very small portion of the metagenome, and therefore the possibility that the high percentage of defense genes is biased toward the low gene count for these phyla cannot be disregarded.

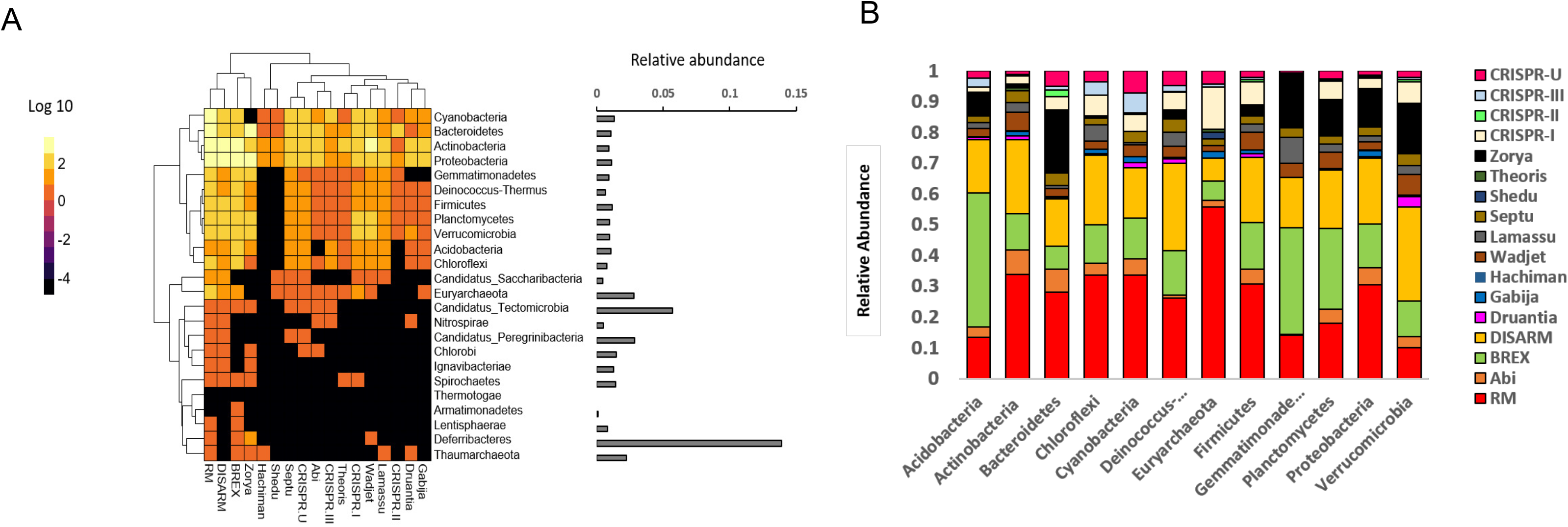

Analysis of the relative contribution of each defense system within each phylum also showed that genes belonging to the RM, DISARM, and BREX systems were the main contributors across the majority of phyla (Figure 1B). The recently discovered Zorya system was predominantly represented in the phyla Gemmatimonadetes, Bacteroidetes, Planctomycetes, Proteobacteria and Verrumicrobia, while CRISPR systems showed the highest contribution in Cyanobacteria and Euryarchaeota. Interestingly, non-canonical anti-phage systems represented more than 50% of the defense systems identified for all phyla aside from Euryarchaeota, with Verrucomicrobia, Planctomycetes and Acidobacteria possessing the highest distribution of non-canonical defense genes.

### Innate immunity is dominated by BREX and DISARM genes

As highlighted above, anti-phage systems across phyla in the hypolith metagenome were dominated by non-canonical innate systems. Further analysis of the distribution of defense genes revealed that anti-phage systems in the majority of phyla were dominated by BREX and DISARM genes. The two systems together accounted for 33.4% of defense genes, compared to 31.7% genes belonging to canonical RM systems.

A total of 3758 genes for the DISARM system were identified. These included the Class I marker gene *drmD* (449 counts, 11.9% of DISARM genes), which encodes the SNF2-like helicase (23), as well as the Class II marker gene *drmA* (1020, 17.1% of DISARM genes), which encodes a protein with a putative helicase domain (23). Similarly, a total of 4598 genes representing all BREX types were identified in the metagenome. Interestingly, the most abundant gene from this system found in the metagenome, *pglW* (2640, 57.4% of BREX genes), which codes for a serine/threonine kinase, is specific to the type 2 BREX system, also called the Pgl system (21). By comparison, of the 7908 RM genes found in the metagenome, the most abundant is hsdM (1423, 18% of RM genes), a type I DNA methylase responsible for the protection of host DNA (24). In fact, more than 50% of RM defense genes were attributed to type I RM systems.

The third non-canonical system representing more than 10% of the anti-phage defense systems in a subset of the phyla, the Zorya system, included a total of 2411 genes in the metagenome. The majority of these were homologous to the two genes that make up a proton channel, *zorA* and *zor*B. This is a common feature in all types of Zorya system and is thought to cause depolarization of the membrane upon infection (22).

### Type I CRISPR-Cas genes comprise the bulk of anti-phage adaptive immunity genes

In total, 2234 CRISPR-*cas* genes were identified in 1601 contigs by searching for shared sequence similarities against the CDD database. A substantial proportion of all classified CRISPR-cas loci (71.4%) belonged to type I CRISPR-Cas systems, followed by type III (18.5%) and type II (10.2%) (Table S5). While the abundance of Cas I-B loci sequences in the public databases suggests that the Cas-I mechanism is the most common in both bacteria and archaea (20 and 30% of total CRISPR loci (25), less than 3% of these loci were present in our composite metagenome (Table S5, Figure 2). Surprisingly, CRISPR-cas loci linked to Types I-C and I-E were the most prevalent, at 24.1% and 12.9% of classified CRISPR-*cas* loci, respectively. Another subtype identified at higher relative abundances than previously reported (25) was I-U, at 10.76% of classified *cas* loci. This subtype is characterized by the marker GSU0054 domain, which was the fourth most abundant cas CDD overall (108 occurrences) after cas4, cas1, and cas2.

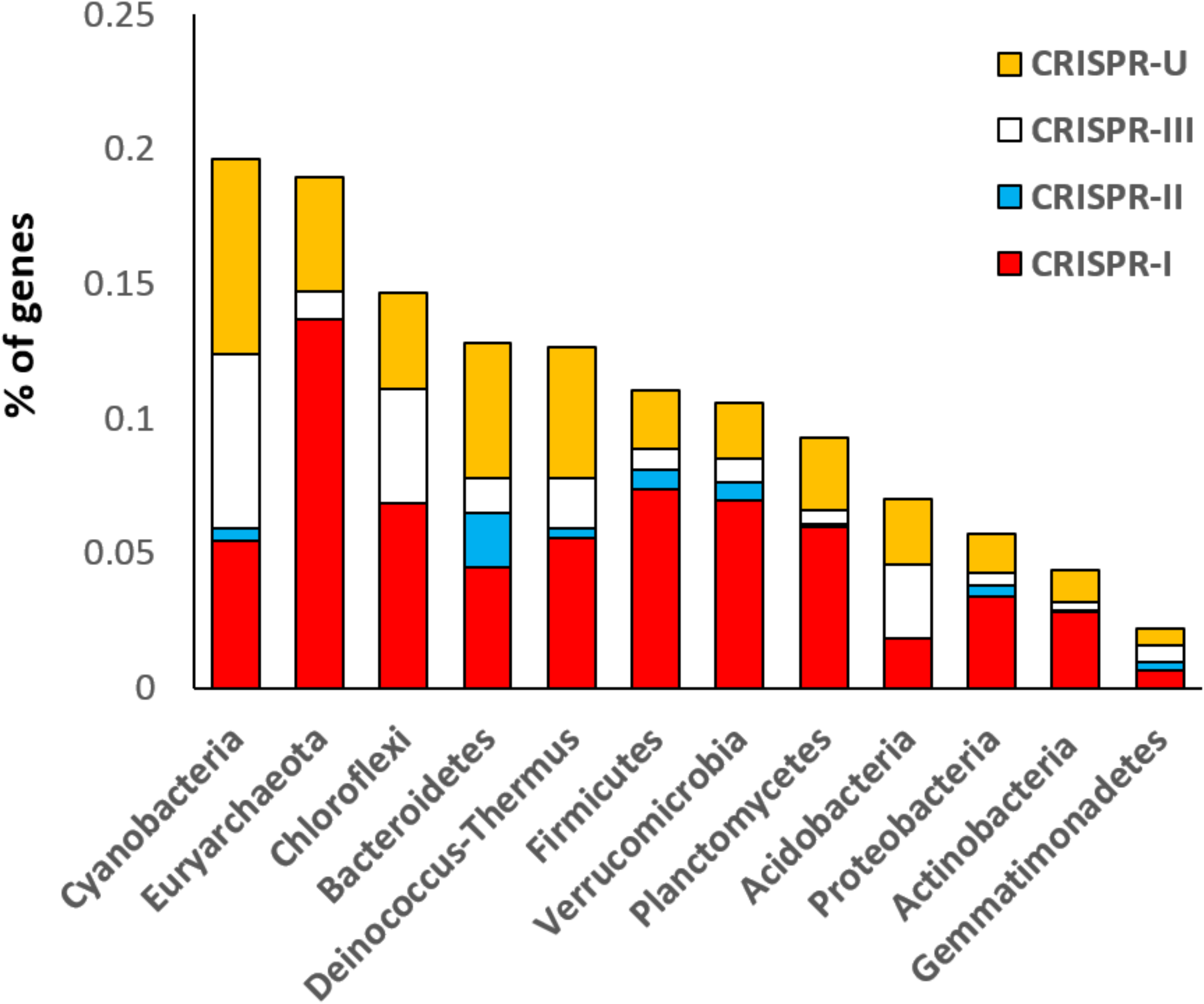

### Phage presence in the niche community is correlated with the CRISPR arrays

CRISPR arrays represent the history of infection by invading DNA (e.g. phages, plasmids (26, 27), and a study of their composition and frequencies provides insights into phage-host interactions in an ecological context (28). A total of 878 CRISPR arrays harboring 10,292 spacers were identified in the metagenome, with an average length of 36 protospacers per array (Figure S1A). CRISPR array sizes ranged from 2 to 249, with the majority (83.5% of total arrays) falling between 2 and 18 protospacers per array (Figure S1B).

The distribution of CRISPR array sizes in the metagenome was compared to data collected from a ground-water microbiome (29), to compare the array size distributions from environments with potentially different phage-host dynamics (11). The results show that CRISPR arrays in the hypolith metagenome exhibited a smaller and narrower size range, compared to the ground-water community metagenome (Figure 3). This suggests the existence of distinct phage infection frequencies between the different environments; i.e., lower infection frequencies in the cold edaphic community.

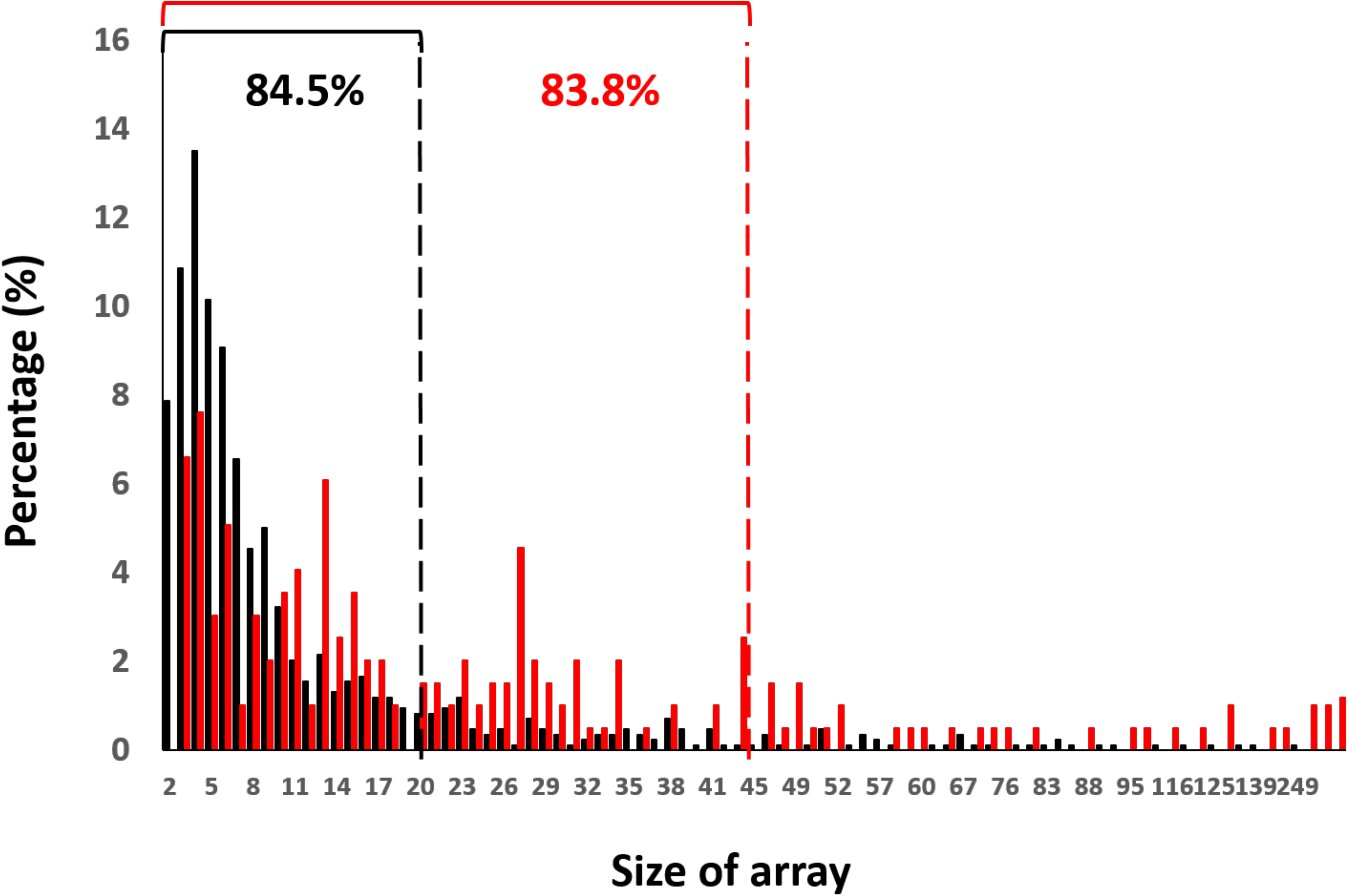

In addition to using the CRISPR array as a tool for understanding infection history, the viral population in the Antarctic soil community was also assessed by assembly of the metavirome. A total of 793 contigs was assembled from the metagenomic sequence data using VirSorter (30). Taxonomic annotation of these contigs, using a database of viral reference genomes (31), unambiguously assigned 645 of these as viral, 560 of which were further assigned to the order of tailed phages *Caudovirales.* Within this order, the majority of contigs were assigned to Siphoviridae (52%), followed by unclassified Caudovirales (14%), and viruses with no assigned family (13%) (Figure S2). To access the correlation between the viral contigs and the CRISPR arrays, spacers from the metagenome were matched to both the VirSorter contigs and a set of contigs from environmental datasets (IMG/VR (32), which allowed for the taxonomic assignment of 394 (3.8% of total number of spacers) CRISPR-cas spacers (Figure S3). The resulting similarity network (Figure 4) showed that all 73 VirSorter phage contigs included in the network (red nodes) matched to CRISPR-cas spacers (grey nodes), suggesting that a substantial fraction (11.3%) of the identified viral population had a history of infection *in situ* in the host population, and may therefore be actively involved in shaping the adaptive immunity of the microbial community. In addition, several distinct clusters showed matches between a single VirSorter contig and several spacers, suggesting these viral contigs are common infection agents.

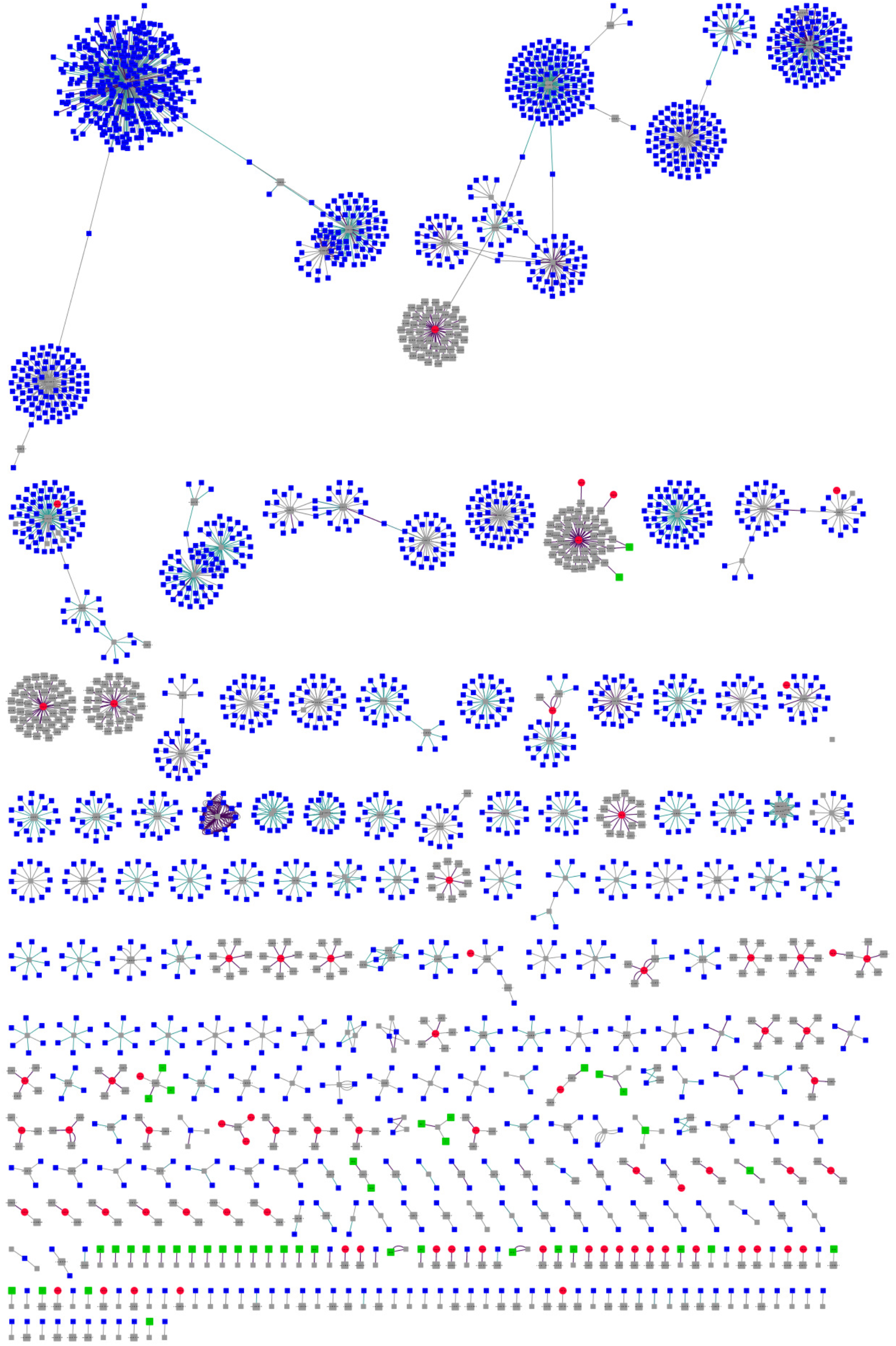

Functional analysis using eggNOG showed the presence of genes that facilitate infection such as genes that code for chitinases, which are involved in the degradation of the protective biofilm (33), as well as a AntA/AntB antirepressor gene, thought to be involved in phage anti-immunity (34) (Figure 5). In addition, the eggNOG functional analysis of the 645 VirSorter viral contigs also revealed the presence of genes contributing to phage virulence (Table S2), the most abundant of which encode for methyltransferases, which are actively involved in the evasion of the R.M systems (35). This result suggests the possibility of an evolutionary pressure for the phages to develop evasion mechanisms against their hosts, which further hints at active phage-host dynamics in these long-enduring Antarctic hypoliths.

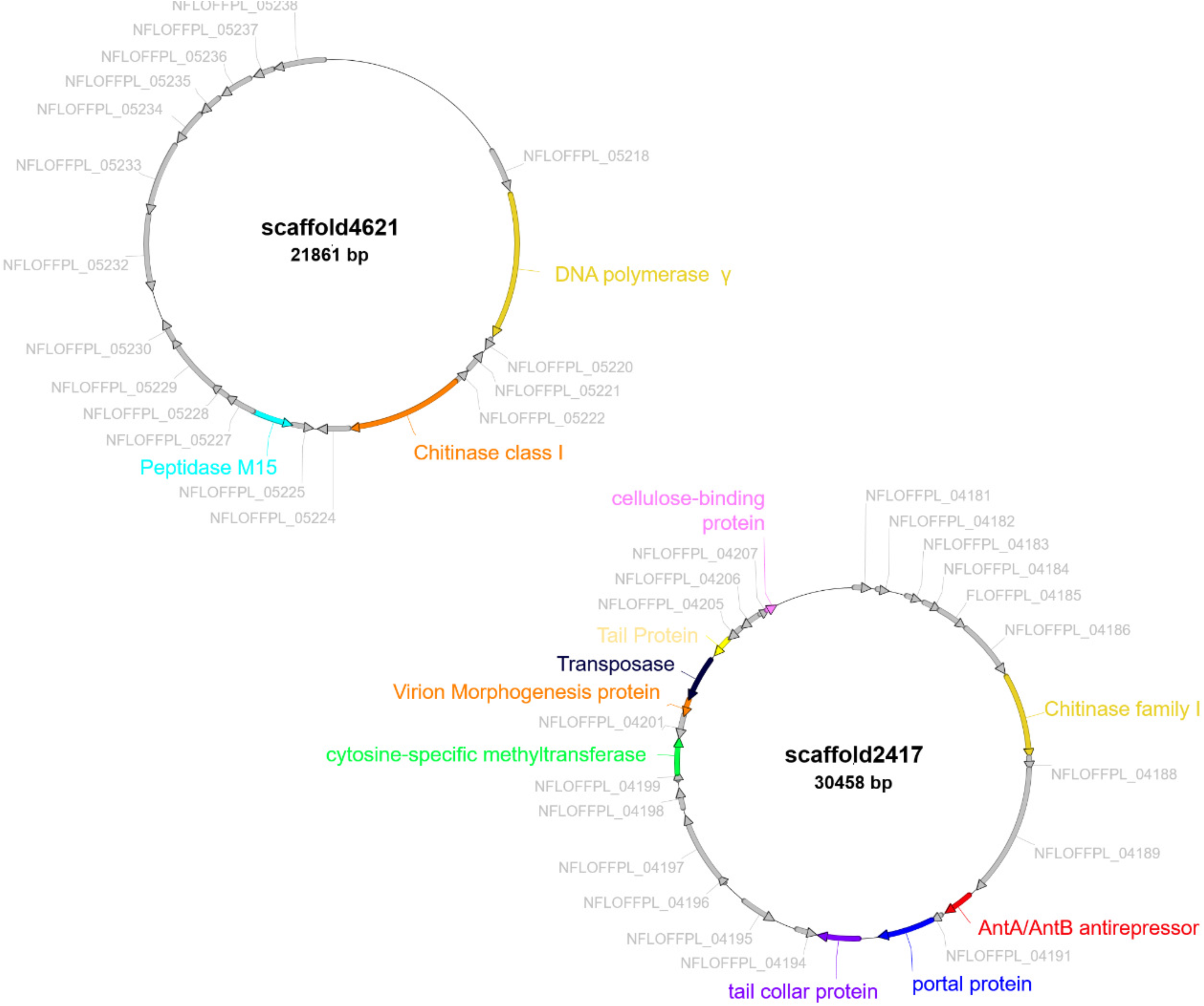

## Discussion

Due to the relatively simple trophic structures in cold desert systems, including Antarctic soils, cryptic microbial communities are considered to be important drivers of local ecosystem services (36). However, the extent to which these communities are influenced by phages remains largely unexplored. Such interactions may shape the diversification and community interactions in cold desert systems. Qualitative surveys of Antarctic metaviromics have reported a high diversity of viruses associated with microbial communities of open soils, and cryptic niches (12, 13). Evidence, albeit limited, that Antarctic soil phages exist predominantly in a lysogenic rather than lytic lifestyle (14), has led to suggestions that the functional role of phages in this spatially restricted, water-constrained desert soil niche may be limited (11).

The results presented in this study provide the first evidence of interaction between phage and hosts in this psychrophilic edaphic environment. This is most evident in the correlation between the metavirome of the hypolith community and the CRISPR-arrays, which suggest the active evolution of the adaptive immune system against local viral threats. This idea of community adaption to local phage threat is further implied by the positive correlation between the CRISPR arrays and viruses extracted from local soils. In fact, a previous study (37) has already suggested that recruitment from surrounding soils plays an important role in the development of hypoliths, and this might also be extended to the recruitment of phages from the surrounding ecosystem. Another indication of active interaction between phage and host is suggested by the presence of several methyltransferases in the metagenome-assembled viral contigs, which are a hallmark of viral evasion against native host RM systems (31, 35). Other genes found in this virome contigs include genes specifically involved in the degradation of biofilm matrices and evasion against RM systems, further suggesting that there is a complex network of interactions at play between phages and their hosts in the hypolithic environment.

While the metagenomics data analysed in this study does not give a direct indication of the temporal scale of the phage-host interactions occurring in the hypolith, the short sizes of CRISPR-array sizes in the hypolith metagenome suggest a low frequency of infection. This low frequency is further hinted at when comparing the hypolith CRISPR-array sizes with those of a more fluid and homogenous environment, where viral-host interactions are assumed to be a frequent occurrence (29). Together, these results imply a model for viral-host interactions in hypoliths that follows the ‘static-step-static’ development model suggested by Pointing et al. (38), driven by the stochastic and intermittent nature of rain events in such water-limited ecosystems. A surprising result from this study is the prevalence of non-canonical innate immunity systems, the most prominent of which are the BREX and DISARM systems. While these two systems have been shown to be widespread in bacteria using a pan-genomic dataset (21, 23), the present study represent the first evidence for the prevalence of these systems in ecological samples. As such, this result implies that non-canonical innate immunity is more important for anti-phage microbial community defense than previously thought and should therefore be the focus for future studies into innate immunity in the ecological context. There are also indications from the hypolith metagenome that the prevalence of non-canonical innate immunity over traditional RM and Abi system for defense against phages is related to the adaptation of the hypolith communities to specific local viral populations. For instance, the Zorya system, the third most prevalent non-canonical immunity system in the metagenome, is hypothesized to operate similarly to the Abi system (22). In turn, Zorya systems provide resistance against a limited range of phages, including the ssDNA family *Microviridae* (22), which has been shown to be prevalent in Antarctic aquatic and soil niches (39).

## Conclusion

Together, these results are not consistent with the suggestion that the constraints of the environment, such as low temperatures, low aw and resulting very limited capacity for inter-particle diffusion, lead to extremely localized phage-host interactions (11). Rather, the data are suggestive of a dynamic and continual interaction between host and phage. Nevertheless, inter-particle communication and exchange may be limited to brief periods when bulk liquid water is present, after snow melt, for example. Furthermore, the low metabolic rates (the inevitable consequence of Arrhenius effects (temperature dependence of reaction rates) in cold environments) should also limit the rates at which phages can replicate and propagate, further limiting the frequency of interactions with their hosts (40). We suggest that the localized nature of host-phage interactions in the hypolith niche and the limited inter-particle communication, where bacterial hosts are not frequently challenged by novel phage threats, leads to a reliance of microbial communities on innate immunity as the primary defense against phage infection. The smaller sizes of CRISPR arrays in the Antarctic soil metagenome sequences compared to those from a temperate aquatic environment, and the under-representation of CRISPR systems, give further credence to the temporally sporadic interaction between phages and their hosts. Nevertheless, the correlation between the metavirome and the CRIPR-cas arrays, together with the presence of bacteriophage evasion genes in the metavirome, suggest that phage-host interactions within the hypolith community are a dynamic process that leads to co-evolution of both phages and hosts. We therefore suggest that phages play a hitherto underestimated role in driving the evolution of Antarctic soil microbial communities by shaping their collective immunity.

## Materials and Methods

### Sample collection, DNA extraction and metagenomic sequencing

The sample collection, DNA extraction and metagenomic sequencing protocols used in this study have been described previously (41). Briefly, a total of 50 samples were collected from hypolithic niches in the Antarctica Miers Valley (GPS 78°09’36.0”S 164°06’00.0”E) and stored in sterile Whirl-Pak bags (Nasco International, Fort Atkinson, WI, USA) at −20 °C. Metagenomic DNA was extracted from each sample using a PowerSoil DNA isolation kit (MO BIO, Carlsbad, CA, USA), and the purified DNA was pooled before further processing. Purified DNA was sheared into fragments of approximately 300 bps and further purified from 1% agarose gels. Subsequent sequencing was performed using Illumina HiSeq-2000 paired-end technology (2 x 101 bp), and the resulting reads were trimmed and assembled as described below.

### Metagenome assembly and taxonomical annotation

Metagenomic DNA sequence data were quality-filtered by trimmomatic version 0.36 using a phred cut-off > 30 (42). The assembly of high-quality reads from the metagenome sequence dataset was conducted using the IDBA-UD tool (43) and contig lengths were extended (scaffolded) using SSPACE Basic (43). The statistics for the assembly of the metagenome are presented in Table S1. Contigs were taxonomically assigned using the MEGAN v6 pipeline (44) with the NCBI taxonomy database for taxon ID assignment.

### Detection of the innate and adaptive defense systems

Metagenomic contigs were used for functional gene predictions using prodigal v2.50, with the –meta parameter implementation (45). Predicted genes were subsequently screened for domain similarity with known defense systems against the conserved domain database (CDD) of clusters of orthologous groups (COGs) and protein families (Pfams) using rps-blast (E value < 1e-02) (33). These results were manually filtered for the identification of phage-specific defense systems, which include restriction-modification (R.M), bacteriophage exclusion (BREX), abortive infection (Abi), defense island system associated with restriction-modification (DISARM), and other recently identified systems using a refined list of COG and Pfam position-specific score matrices (PSSMs) for marker genes in these systems (21-23, 46). A list of the marker genes used in this study can be found in Table S2. Additionally, defense genes that could not be clustered into a specific system were classified as ambiguous as were not considered for subsequent analysis (Table S3).

ORFs predicted using prodigal v2.50 were queried against the CDD database for the presence of putative CRISPR-cas genes (47), using delta-blast at a cutoff E value of 1e-03. Multi-gene cas modules were identified as those having multiple cas annotated genes with ≤5 ORF spacings. Type and subtype classifications were assigned following the updated classification set by Makarova et al. (25).

### Phage genome identification and CRISPR spacer matching

Antarctic hypolith phage genomes were identified from the assembled metagenome using VirSorter (30) on the iVirus platform hosted by Cyverse (48), using the virome database and the microbial decontamination option. Only predictions of categories 1, 2, 4 and 5 were used (phages and prophages identified with the “pretty sure” and “quite sure” qualification). Additional phage environmental phage contigs were downloaded from the IMG/VR database version 2018-07-01_4 (32) and used for the network construction. Taxonomic assignment of assembled contigs was performed by using the DIAMOND blastx function with a viral database downloaded from the NCBI Viral Genomes Resource and e-value set to 1e-5. ORFs of VirSorter contigs were predicted using Prodigal v2.50 (31, 49) with the virus genomes setting and annotated using eggNOG-mapper v1 (50) with the DIAMOND option and the EggNOG v4.5.1 database (51). Annotation were visualized with the ApE v2.0.55 plasmid editor (http://jorgensen.biology.utah.edu/wayned/ape/).

The CRISPR recognition tool (CRT) v1.2 was used with the default settings to search for CRISPR arrays in the hypolith metagenome (52). The identified spacers in the arrays were matched with the VirSorter phage database and the IMG/VR database using blastn of the BLAST+ suite with the following parameters: - qcov_hsp_perc 80 -task blastn -dust no -soft_masking false (53). Spacer matches of > 90% sequence identity for the VirSorter genomes and >95% identity for the IMG/VR genomes were exported and visualized as a network in Cytoscape (54), where the nodes are spacers (grey) or genomes (blue = IMG/VR; red = VirSorter) and the edges blastn matches.

## Acknowledgements

We are grateful to the National Research Foundation (NRF) (Grant ID 118981, the South African National Antarctic Programme (SANAP 110717), and the University of Pretoria for funding. TPM also wishes to acknowledge the Fulbright Visiting Scholar Program for providing sabbatical funding. EMA gratefully acknowledges the support of the Biotechnology and Biological Sciences Research Council (BBSRC); this research was funded by the BBSRC Institute Strategic Programme Gut Microbes and Health BB/R012490/1 and its constituent project(s) BBS/E/F/000PR10353 and BBS/E/F/000PR10356

## References

1. Makhalanyane TP, Van Goethem MW, Cowan DA. 2016. Microbial diversity and functional capacity in polar soils. Current Opinion in Biotechnology 38:159–166.

2. Pearce DA. 2012. Extremophiles in Antarctica: Life at low temperatures, p 87–118. *In* Stan-Lotter H, Fendrihan S (ed), Adaption of Microbial Life to Environmental Extremes. Springer Vienna.

3. Casanueva A, Tuffin M, Cary C, Cowan DA. 2010. Molecular adaptations to psychrophily: the impact of ‘omic’ technologies. Trends in microbiology 18:374–81.

4. de Scally SZ, Makhalanyane TP, Frossard A, Hogg ID, Cowan DA. 2016. Antarctic microbial communities are functionally redundant, adapted and resistant to short term temperature perturbations. Soil Biology & Biochemistry 103:160–170.

5. de los Rios A, Wierzchos J, Sancho LG, Ascaso C. 2004. Exploring the physiological state of continental Antarctic endolithic microorganisms by microscopy. FEMS microbiology ecology 50:143–52.

6. Yergeau E, Kowalchuk GA. 2008. Responses of Antarctic soil microbial communities and associated functions to temperature and freeze-thaw cycle frequency. Environmental microbiology 10:2223–35.

7. Cowan DA, Makhalanyane TP, Dennis PG, Hopkins DW. 2014. Microbial ecology and biogeochemistry of continental Antarctic soils. Frontiers in Microbiology 5:154.

8. Hogg ID, Cary SC, Convey P, Newsham KK, O’Donnell AG, Adams BJ, Aislabie J, Frati F, Stevens MI, Wall DH. 2006. Biotic interactions in Antarctic terrestrial ecosystems: Are they a factor? Soil biology & biochemistry 38:3035–3040.

9. Caruso T, Hogg ID, Nielsen UN, Bottos EM, Lee CK, Hopkins DW, Cary SC, Barrett JE, Green TA, Storey BC. 2019. Nematodes in a polar desert reveal the relative role of biotic interactions in the coexistence of soil animals. Communications biology 2:63.

10. Lambrechts S, Willems A, Tahon G. 2019. Uncovering the uncultivated majority in antarctic soils: toward a synergistic approach. Frontiers in microbiology 10:242.

11. Zablocki O, Adriaenssens EM, Cowan D. 2016. Diversity and ecology of viruses in hyperarid desert soils. Appl Environ Microbiol 82:770–777.

12. Adriaenssens EM, Kramer R, Van Goethem MW, Makhalanyane TP, Hogg I, Cowan DA. 2017. Environmental drivers of viral community composition in Antarctic soils identified by viromics. Microbiome 5.

13. Zablocki O, van Zyl L, Adriaenssens EM, Rubagotti E, Tuffin M, Cary SC, Cowan D. 2014. High-level diversity of tailed phages, eukaryote-associated viruses, and virophage-like elements in the metaviromes of antarctic soils. Appl Environ Microbiol 80:6888–6897.

14. Williamson KE, Radosevich M, Smith DW, Wommack KE. 2007. Incidence of lysogeny within temperate and extreme soil environments. Environmental microbiology 9:2563–2574.

15. Wei ST, Lacap-Bugler DC, Lau MC, Caruso T, Rao S, de los Rios A, Archer SK, Chiu JM, Higgins C, Van Nostrand JD. 2016. Taxonomic and functional diversity of soil and hypolithic microbial communities in Miers Valley, McMurdo Dry Valleys, Antarctica. Frontiers in microbiology 7:1642.

16. Wei ST, Higgins CM, Adriaenssens EM, Cowan DA, Pointing SB. 2015. Genetic signatures indicate widespread antibiotic resistance and phage infection in microbial communities of the McMurdo Dry Valleys, East Antarctica. Polar Biology 38:919–925.

17. Gómez P, Buckling A. 2011. Bacteria-phage antagonistic coevolution in soil. Science 332:106–109.

18. Stern A, Sorek R. 2011. The phage - host arms race: shaping the evolution of microbes. Bioessays 33:43–51.

19. Koskella B, Brockhurst MA. 2014. Bacteria–phage coevolution as a driver of ecological and evolutionary processes in microbial communities. FEMS microbiology reviews 38:916–931.

20. Makarova KS, Wolf YI, Koonin EV. 2013. Comparative genomics of defense systems in archaea and bacteria. Nucleic acids research 41:4360–4377.

21. Goldfarb T, Sberro H, Weinstock E, Cohen O, Doron S, Charpak-Amikam Y, Afik S, Ofir G, Sorek R. 2015. BREX is a novel phage resistance system widespread in microbial genomes. The EMBO journal 34:169–183.

22. Doron S, Melamed S, Ofir G, Leavitt A, Lopatina A, Keren M, Amitai G, Sorek R. 2018. Systematic discovery of antiphage defense systems in the microbial pangenome. Science 359:eaar4120.

23. Ofir G, Melamed S, Sberro H, Mukamel Z, Silverman S, Yaakov G, Doron S, Sorek R. 2018. DISARM is a widespread bacterial defence system with broad anti-phage activities. Nature microbiology 3:90.

24. Loenen WA, Dryden DT, Raleigh EA, Wilson GG. 2013. Type I restriction enzymes and their relatives. Nucleic acids research 42:20–44.

25. Makarova KS, Wolf YI, Alkhnbashi OS, Costa F, Shah SA, Saunders SJ, Barrangou R, Brouns SJ, Charpentier E, Haft DH. 2015. An updated evolutionary classification of CRISPR–Cas systems. Nature Reviews Microbiology 13:722.

26. Datsenko KA, Pougach K, Tikhonov A, Wanner BL, Severinov K, Semenova E. 2012. Molecular memory of prior infections activates the CRISPR/Cas adaptive bacterial immunity system. Nature communications 3:945.

27. Fineran PC, Charpentier E. 2012. Memory of viral infections by CRISPR-Cas adaptive immune systems: acquisition of new information. Virology 434:202–209.

28. Vale PF, Little TJ. 2010. CRISPR-mediated phage resistance and the ghost of coevolution past. Proceedings of the Royal Society B: Biological Sciences 277:2097–2103.

29. Burstein D, Sun CL, Brown CT, Sharon I, Anantharaman K, Probst AJ, Thomas BC, Banfield JF. 2016. Major bacterial lineages are essentially devoid of CRISPR-Cas viral defence systems. Nature communications 7:10613.

30. Roux S, Enault F, Hurwitz BL, Sullivan MB. 2015. VirSorter: mining viral signal from microbial genomic data. PeerJ 3:e985.

31. Brister JR, Ako-Adjei D, Bao Y, Blinkova O. 2014. NCBI viral genomes resource. Nucleic acids research 43:D571–D577.

32. Paez-Espino D, Chen I-MA, Palaniappan K, Ratner A, Chu K, Szeto E, Pillay M, Huang J, Markowitz VM, Nielsen T. 2016. IMG/VR: a database of cultured and uncultured DNA Viruses and retroviruses. Nucleic acids research:gkw1030.

33. Altschul SF, Gish W, Miller W, Myers EW, Lipman DJ. 1990. Basic local alignment search tool. Journal of molecular biology 215:403–410.

34. Ravin NV, Svarchevsky AN, Dehò G. 1999. The anti - immunity system of phage - plasmid N15: identification of the antirepressor gene and its control by a small processed RNA. Molecular microbiology 34:980–994.

35. Labrie SJ, Samson JE, Moineau S. 2010. Bacteriophage resistance mechanisms. Nature Reviews Microbiology 8:317.

36. Cary SC, McDonald IR, Barrett JE, Cowan DA. 2010. On the rocks: the microbiology of Antarctic Dry Valley soils. Nature Reviews Microbiology 8:129–38.

37. Makhalanyane TP, Valverde A, Lacap DC, Pointing SB, Tuffin MI, Cowan DA. 2013. Evidence of species recruitment and development of hot desert hypolithic communities. Environmental Microbiology Reports 5:219–224.

38. Pointing SB, Warren-Rhodes KA, Lacap DC, Rhodes KL, McKay CP. 2007. Hypolithic community shifts occur as a result of liquid water availability along environmental gradients in China’s hot and cold hyperarid deserts. Environmental microbiology 9:414–24.

39. Daniel AdC, Pedrós-Alió C, Pearce DA, Alcamí A. 2016. Composition and interactions among bacterial, microeukaryotic, and T4-like viral assemblages in lakes from both polar zones. Frontiers in microbiology 7:337.

40. D’Amico S, Collins T, Marx JC, Feller G, Gerday C. 2006. Psychrophilic microorganisms: challenges for life. EMBO Rep 7:385–389.

41. Guerrero LD, Vikram S, Makhalanyane TP, Cowan DA. 2017. Evidence of microbial rhodopsins in Antarctic Dry Valley edaphic systems. Environmental Microbiology 19:3755–3767.

42. Bolger AM, Lohse M, Usadel B. 2014. Trimmomatic: a flexible trimmer for Illumina sequence data. Bioinformatics 30:2114–2120.

43. Peng Y, Leung HC, Yiu S-M, Chin FY. 2012. IDBA-UD: a de novo assembler for single-cell and metagenomic sequencing data with highly uneven depth. Bioinformatics 28:1420–1428.

44. Boetzer M, Henkel CV, Jansen HJ, Butler D, Pirovano W. 2010. Scaffolding preassembled contigs using SSPACE. Bioinformatics 27:578–579.

45. Huson DH, Auch AF, Qi J, Schuster SC. 2007. MEGAN analysis of metagenomic data. Genome research 17:377–386.

46. Makarova KS, Haft DH, Barrangou R, Brouns SJ, Charpentier E, Horvath P, Moineau S, Mojica FJ, Wolf YI, Yakunin AF. 2011. Evolution and classification of the CRISPR–Cas systems. Nature Reviews Microbiology 9:467.

47. Marchler-Bauer A, Bo Y, Han L, He J, Lanczycki CJ, Lu S, Chitsaz F, Derbyshire MK, Geer RC, Gonzales NR. 2016. CDD/SPARCLE: functional classification of proteins via subfamily domain architectures. Nucleic acids research 45:D200–D203.

48. Bolduc B, Youens-Clark K, Roux S, Hurwitz BL, Sullivan MB. 2017. iVirus: facilitating new insights in viral ecology with software and community data sets imbedded in a cyberinfrastructure. The ISME journal 11:7.

49. Hyatt D, Chen G-L, LoCascio PF, Land ML, Larimer FW, Hauser LJ. 2010. Prodigal: prokaryotic gene recognition and translation initiation site identification. BMC bioinformatics 11:119.

50. Huerta-Cepas J, Forslund K, Coelho LP, Szklarczyk D, Jensen LJ, von Mering C, Bork P. 2017. Fast genome-wide functional annotation through orthology assignment by eggNOG-mapper. Molecular biology and evolution 34:2115–2122.

51. Huerta-Cepas J, Szklarczyk D, Forslund K, Cook H, Heller D, Walter MC, Rattei T, Mende DR, Sunagawa S, Kuhn M. 2015. eggNOG 4.5: a hierarchical orthology framework with improved functional annotations for eukaryotic, prokaryotic and viral sequences. Nucleic acids research 44:D286–D293.

52. Bland C, Ramsey TL, Sabree F, Lowe M, Brown K, Kyrpides NC, Hugenholtz P. 2007. CRISPR recognition tool (CRT): a tool for automatic detection of clustered regularly interspaced palindromic repeats. BMC bioinformatics 8:209.

53. Camacho C, Coulouris G, Avagyan V, Ma N, Papadopoulos J, Bealer K, Madden TL. 2009. BLAST+: architecture and applications. BMC bioinformatics 10:421.

54. Shannon P, Markiel A, Ozier O, Baliga NS, Wang JT, Ramage D, Amin N, Schwikowski B, Ideker T. 2003. Cytoscape: a software environment for integrated models of biomolecular interaction networks. Genome research 13:2498–2504.

